# Enhancer transcription identifies *cis*-regulatory elements for photoreceptor cell types

**DOI:** 10.1101/513598

**Authors:** Rangarajan D. Nadadur, Carlos Perez-Cervantes, Nicolas Lonfat, Linsin A. Smith, Andrew E. O. Hughes, Sui Wang, Joseph C. Corbo, Connie Cepko, Ivan P. Moskowitz

## Abstract

Identification of the *cis*-regulatory elements (CREs) that regulate gene expression in specific cell types is critical for defining the gene regulatory networks (GRNs) that control normal physiology and disease states. We previously utilized non-coding RNA (ncRNA) profiling to define CREs that comprise a GRN in the adult mouse heart^1^. Here, we applied ncRNA profiling to the mouse retina in the presence and absence of *Nrl*, a rod photoreceptor-specific transcription factor required for rod versus cone photoreceptor cell fate. Differential expression of *Nrl*-dependent ncRNAs positively correlated with differential expression of *Nrl*-dependent local genes. Two distinct *Nrl*-dependent regulatory networks were discerned in parallel: *Nrl*-activated ncRNAs were enriched for accessible chromatin in rods but not cones whereas *Nrl*-repressed ncRNAs were enriched for accessible chromatin in cones but not rods. Furthermore, differential *Nrl*-dependent ncRNA expression levels quantitatively correlated with photoreceptor cell type-specific ATAC-seq read density. Direct assessment of *Nrl*-dependent ncRNA-defined loci identified functional cone photoreceptor CREs. This work supports differential ncRNA profiling as a platform for identifying context-specific regulatory elements and provides insight into the networks that define photoreceptor cell types.

## Introduction

Identification of tissue and context specific *cis*-regulatory elements (CREs) is critical to defining the transcriptional networks that govern physiology and disease across biological contexts. The advent of high-throughput sequencing has allowed for genome-wide analysis of chromatin state as a proxy for regulatory elements^2^. The use of histone modifications, chromatin status and transcription factor (TF) occupancy to define gene regulation has been successful in many contexts^3–6^. However, these approaches lack both specificity and quantitative resolution of enhancer activity^2^. These observations indicate the need for complementary strategies for the identification of functional context-dependent enhancers.

A growing body of literature indicates that noncoding RNAs (ncRNAs) are transcribed from active CREs. The function of these transcripts and their role in gene regulation is an area of active research^1,7–10^. We previously demonstrated that differential enhancer transcription can be used to define a gene regulatory network (GRN)^1^. We identified ncRNAs whose expression was dependent on the cardiac transcription factor TBX5^1^. These genome wide TBX5-dependent ncRNAs from the adult mouse atria defined a GRN for cardiac rhythm^1^. This approach led to the identification of potent regulatory elements in cardiomyocytes and identified functional ncRNAs that mediated TBX5-dependent gene regulation. These findings suggested that differential enhancer transcription may be an effective complementary approach to chromatin accessibility, epigenetic marks, and reporter assays for the discovery of context-dependent CREs. However, the applicability of this approach across broader contexts and its ability to distinguish activated and repressed elements has not been examined.

Here, we applied the differential ncRNA approach to identify regulatory elements that are active in either of two specific cell types, rod and cone photoreceptors, in the mouse retina. Rod photoreceptors are active in dim light and constitute the most abundant retinal cell type, comprising ~80% of all mouse retinal cells and 95% of human photoreceptors^11,12^. In contrast, cone photoreceptors are active in bright light and mediate high-acuity vision and color vision. Their critical role in daylight vision makes them a desirable cell type to replace from stem cells, or to target for gene therapy, in diseases that lead to blindness^13^. An understanding of the GRNs that control cone versus rod fate is essential for understanding both normal retinal biology and for cone replacement, as has been recently demonstrated for rods in mice^14–19^. In addition, gene therapy vectors that require expression specifically in rods andlor cones would benefit from a broader range of validated photoreceptor CREs^20–22^.

Rods and cones are produced by retinal progenitor cells (RPCs), with cones generally produced earlier in development than rods, from RPCs that express the TFs *Otx2, Olig2*, and *Oc1*^23–26^ These and other TFs with essential roles in RPCs, rods, or cones have been identified^27,28^. In addition to *Otx2*, a close homologue, *Crx*, is required for normal gene expression in both rods and cones^29–31^. In contrast, *Nrl*, a basic leucine zipper TF, is expressed only in rods and is required for their formation. In *Nrl*^-/-^ mutant mice, rod photoreceptors fail to form and are instead transformed into cells that resemble cone photoreceptors in most respects^32^.

We aimed to identify the photoreceptor CREs that regulate rod and cone gene expression programs using differential ncRNA transcriptome analysis. We compared ncRNA abundance genome-wide in the wild-type versus *Nrl*^-/-^ mouse retina. We found that *Nrl*-activated versus repressed ncRNA transcripts defined photoreceptor regulatory elements. *Nrl*-activated ncRNAs, predominant in the wild-type retina, identified locations of accessible chromatin in rods and that were near rod-expressed genes. In contrast, *Nrl*-repressed ncRNAs, predominant in the *Nrl*^-/-^ retina, identified locations of accessible chromatin in cones that were near cone-expressed genes. Moreover, differential expression of *Nrl*-dependent ncRNAs and that of local target genes were quantitatively correlated. Furthermore, the change in expression of these ncRNAs positively correlated with differential signal from rod and cone ATAC-seq. Direct assessment of *Nrl*-repressed loci identified active elements for photoreceptor expression, enriched for cone-specific genes. These data illustrate the utility of differential ncRNA profiling for nominating TF-dependent and context-specific regulatory elements.

## Results

### *Nrl*-dependent coding and non-coding transcriptional profiling identifies photoreceptor CREs

We interrogated *Nrl*-dependent coding and non-coding transcription in the mouse retina. We performed mRNA transcriptional profiling to determine *Nrl*-dependent coding gene expression (Figure 1A). We sequenced cDNA libraries made from polyA+ selected RNA from retinas of litter-matched WT and *Nrl* mutant adult mice at postnatal day 21 (P21) (*Nrl*^+/+^ vs *Nrl*^-/-^, n=5 and n= 6 resp., Figure 1A). By P21, mouse photoreceptor differentiation is largely complete. Genotype described 98% of the variance between samples, indicating a specific effect mediated by *Nrl* deletion across biological replicates (Figure 1B). Differential expression testing revealed 4,315 misregulated genes (Figure 1A). *Nrl* expression was absent from *Nrl*^-/-^ samples, along with numerous known *Nrl* targets and rod-specific genes, including *Rho, Gnat1*, and *Nr2e3*. Conversely, genes whose expression is normally absent in rods, including those involved in cone differentiation showed activation in *Nrl*^-/-^ samples, including *Gnat2, Gngt2*, and *Opn1sw*^18^. These expression changes were consistent with previously identified *Nrl*-dependent gene expression and rod-versus cone-specific gene expression patterns in the adult retina^16–18,33,34^.

**Figure 1:**
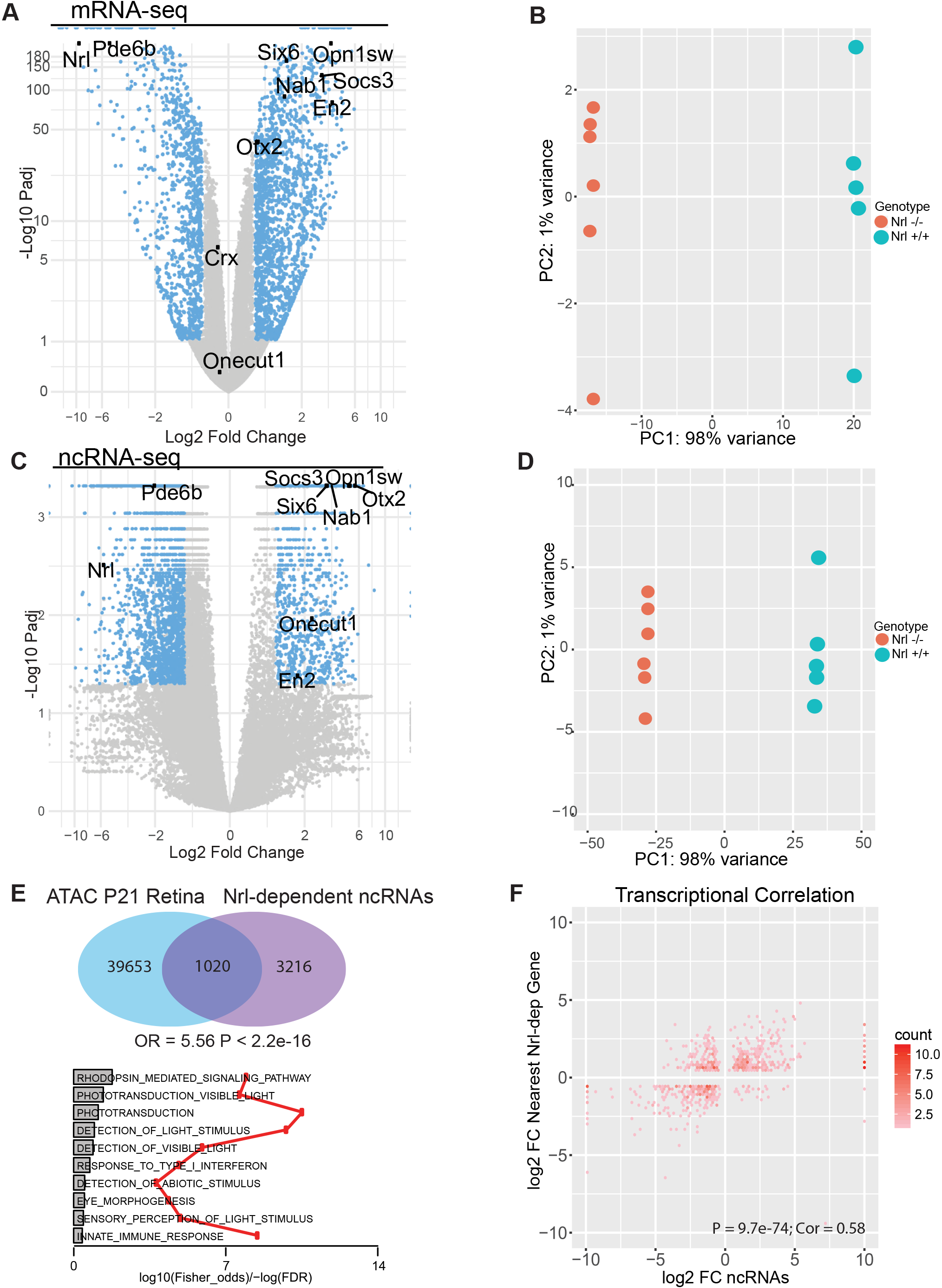
Transcriptional profiling identifies M7-dependent coding and noncoding RNAs. A) Volcano-plot (log_2_ (fold-change) vs -log_10_ (P_Adj_)) of *Nrl*^+/+^ vs *Nrl*^-/-^ coding transcripts. Significantly misregulated genes (P_Adj_ < 0.05) in blue, and non-significant in grey. Selected significant coding genes known to play a role in photoreceptor differentiation labeled (black). B) Principle component analysis on WT and *Nrl*^-/-^ coding transcripts. Samples segregate primarily by genotype, describing 98% of variance on PC1. C) Volcano-plot (log_2_ (fold-change) vs -log_10_ (P_Adj_)) of *Nrl*^+/+^ vs *Nrl*^-/-^ non-coding RNAs (ncRNAs). Significantly misregulated ncRNAs (P_Adj_ < 0.05) are depicted in blue and non-significant in grey. Selected transcripts labeled by nearest *Nrl*-dependent coding genes (black). D) Principle component analysis on WT and *Nrl*^-/-^ noncoding transcripts. Samples segregate primarily by genotype, describing 98% of variance on PC1. D) Top: Venn diagram of the overlap of ATAC-seq from control mouse retina (GSE72550) and *Nrl*-dependent ncRNA transcripts (OR = 5.56, P < 2.2e-16). P-values and Odds Ratios from Fisher Exact Test. Bottom: Gene ontology (GO) analysis of nearby (2Mb) *Nrl*-dependent genes to defined *Nrl*-dependent ncRNAs. Fisher Odds (gray) and -log(FDR) (Red). Significantly enriched GO terms are related to visual processes, including “rhodopsin mediated signaling”, “phototransduction”, and “detection of light stimulus”. F) Scatterplot in hexagonal binning for the *Nrl*-dependent ncRNAs. Differential expression fold change (log2) of nearest *Nrl*-dependent gene vs differential expression fold change (log2) of ncRNAs (x-axis).

We performed ncRNA transcriptional profiling to identify *Nrl*-dependent ncRNAs. We performed deep sequencing of non-polyadenylated RNA from the same control and *Nrl* mutant retinal samples described above (Figure 1C)^35^. This approach identified approximately 30,000 retinal non-coding transcripts by *de novo* transcript assembly (Supplemental FIgure). Approximately 4,657 of these transcripts were *Nrl*-dependent intergenic ncRNAs (FDR<0.05, FC>2). Genotype described 98% of the variance between samples, indicating a specific effect mediated by *Nrl* deletion (Figure 1D). Of the 4,657 *Nrl*-dependent ncRNAs, 3,112 were significantly downregulated, or *Nrl*-activated, and 1,545 were significantly upregulated, or *Nrl*-repressed (Figure 1C).

We hypothesized that some *Nrl*-dependent noncoding transcripts marked *Nrl*-dependent retinal enhancers. Active regulatory elements are characterized by characteristic genomic patterns, including open chromatin^3–5^. To examine the epigenomic landscape of mouse retinal tissue, we analyzed <u>A</u>ssay for <u>T</u>ranspose <u>A</u>ccessible <u>C</u>hromatin (ATAC-Seq) data from whole mouse wild-type retinal tissue at P21 (GSE72550)^19,36,37^. *Nrl*-dependent ncRNAs significantly overlapped with locations of accessible chromatin in the wild-type retina (Figure 1E, top).

We attempted to affiliate *Nrl*-dependent ncRNA-defined regulatory elements with candidate target *Nrl*-dependent coding genes. We sought *Nrl*-dependent mRNAs within 2MB of the ncRNA locus, a conservative consideration of potential distance parameters for CREs^1^. We performed Gene Ontology (GO) analysis of the *Nrl*-dependent genes associated with *Nrl*-dependent ncRNAs. This analysis described enrichment for GO terms related to phototransduction, sensory perception, and response to light stimulus, all consistent with photoreceptor gene expression (Figure 1E, bottom).

We and others have previously hypothesized that quantitative changes in CRE transcription mirrors quantitative changes in CRE activity^1,38^. Identification of CREs using ncRNA transcriptional profiling affords the determination of context-dependent quantitative changes in CRE transcription. We examined the quantitative correlation between the change in *Nrl*-dependent ncRNA transcription and the change in the most proximal *Nrl*-dependent gene. First, we observed that the directionality of the change of each *Nrl*-dependent ncRNA and its local mRNA were significantly concordant. Secondly, the relative expression of *Nrl*-dependent ncRNAs was positively correlated with the relative expression of the most proximal *Nrl*-dependent gene (Figure 1F Cor=0.58, P=9.7e-74). Together, these observations indicate a positive quantitative relationship between enhancer transcription and target gene regulation. Together these observations indicate that *Nrl*-dependent ncRNA profiling identifies genomic regions overlapping with open chromatin and that quantitatively correlate with local *Nrl*-dependent gene expression.

### *Nrl*-dependent ncRNA transcriptional profiling identifies two distinct gene regulatory signatures, specific to rod and cone photoreceptors

We hypothesized that removal of *Nrl* revealed two distinct regulatory networks: a pathway driven by *Nrl* in the *wild-type* retina, composed of genes that require *Nrl* directly or indirectly and are thereby downregulated in *Nrl*^-/-^ samples, referred to as *Nrl*-activated genes; and a pathway that emerged in the *Nrl* mutant retina, composed of genes whose transcription is directly or indirectly negatively regulated by *Nrl* and are thereby upregulated in *Nrl*^-/-^ samples, referred to as *Nrl*-repressed genes. NRL is a well-described driver of rod differentiation and repressor of cone differentiation ^16,32,39–41^. We therefore hypothesized that *Nrl*-activated ncRNAs, higher in the wild-type retina, would specifically define CREs active in rods, whereas the *Nrl*-repressed ncRNAs, higher in the *Nrl* mutant retina, would specifically define CREs active in cones.

We attempted to identify candidate rod and cone CREs by comparing *Nrl*-dependent ncRNAs with open chromatin signatures from rod and cone specific cell populations (GSE83312)^18^, as opposed to whole retinal tissue (Figure 1E top). We found that both *Nrl*-activated and *Nrl*-repressed ncRNAs overlap with open chromatin in both rods and cones (Figure 2A-B, top). However, the *Nrl*-activated ncRNAs showed much greater enrichment for open chromatin in rods relative to open chromatin in cones (Figure 2A-B, OR=2.77 vs 2.36). Conversely, *Nrl*-repressed ncRNAs showed much greater enrichment with cone open chromatin than with rod open chromatin (Figure 2A-B, OR = 5.15 vs 3.35). We interrogated the distribution of activated and repressed *Nrl*-dependent ncRNAs with chromatin accessibility to define rod-specific, cone-specific, and shared CREs. By overlapping rod and cone ATAC-seq sets, we identified total regions of accessible chromatin, of which 25,666 were shared between rods and cones, 9,116 were rod-specific, and 9,148 were cone-specific (Figure 2C). Intersection of *Nrl*-dependent ncRNAs with each group revealed that *Nrl*-activated and *Nrl*-repressed ncRNAs most frequently emanated from regions that were accessible in both rods and cones, suggesting that the majority of *Nrl*-dependent elements are shared photoreceptor elements common to all photoreceptor types (Figure 2C). However, when comparing the pattern of *Nrl*-dependent ncRNA expression from regions accessible in only rods or cones, a cell type-specific pattern emerged: *Nrl*-activated ncRNAs overlapped more frequently with regions only accessible in rods versus cones (105 vs 38, p=7.9e-10, Figure 2C), whereas *Nrl*-repressed ncRNAs overlapped more frequently with regions only accessible in cones versus rods (Figure 2C, 137 vs 10, p < 2.2e-16). These comparisons indicate that *Nrl*-dependent ncRNAs can be separated into three distinct bins of *Nrl*-dependent candidate CREs; *Nrl*-activated or *Nrl*-repressed ncRNAs at shared rod and cone accessible regions; *Nrl*-activated ncRNAs, expressed in the wild-type retina, at regions accessible only in rods, and *Nrl*-repressed ncRNAs, expressed in the *Nrl*^-/-^ retina, at regions accessible only in cones.

**Figure 2:**
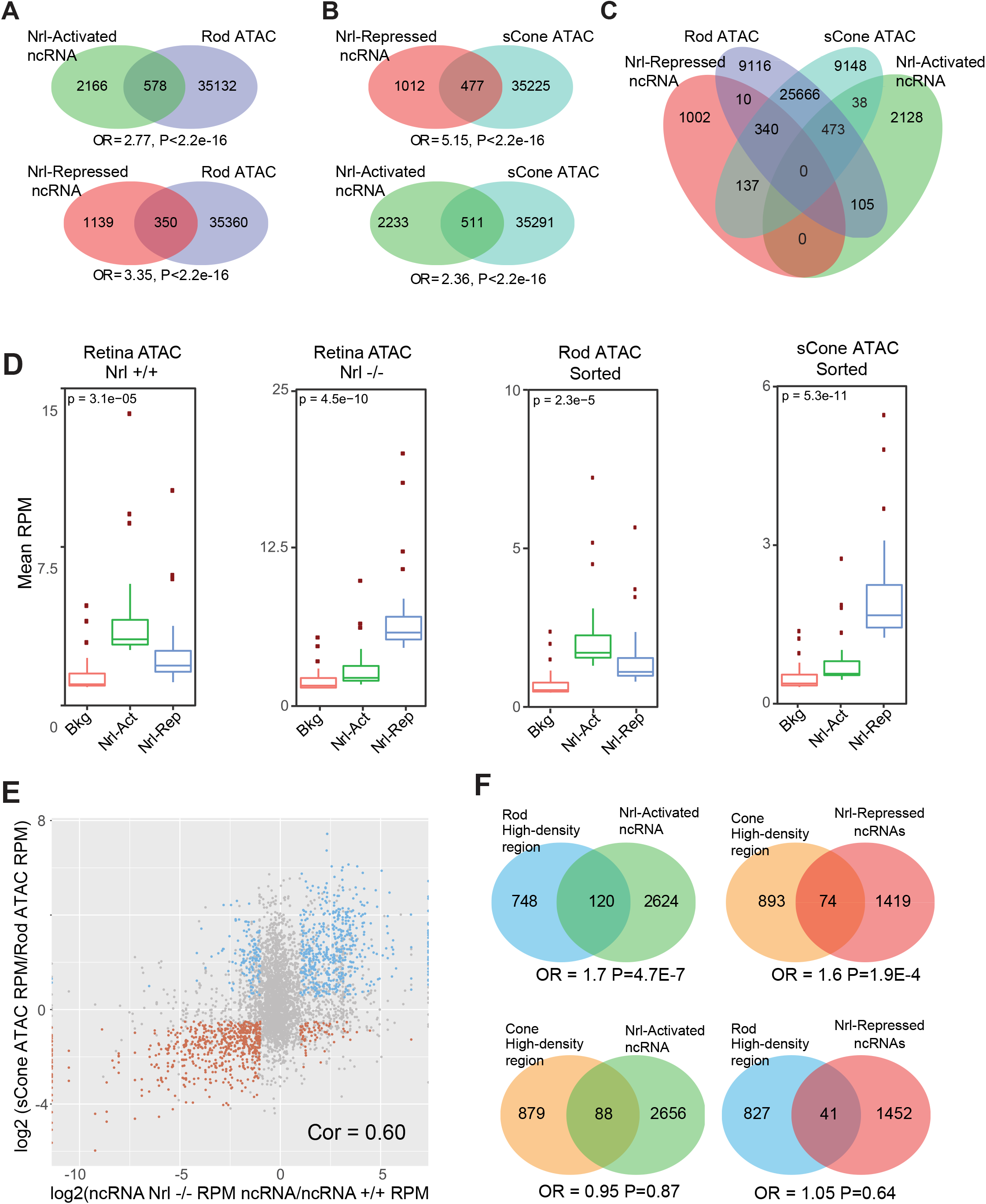
*Nrl*-activated and repressed ncRNAs identify cell-type specific regulatory programs. A) Venn diagrams of the overlap of *Nrl*-activated (top) and *Nrl*-repressed (bottom) ncRNAs with ATAC-seq from rod photoreceptors (GSE83312). P-values and Odds Ratios from Fisher Exact Test. B) Venn diagrams of the overlap of *Nrl*-activated (top) and *Nrl*-repressed (bottom) ncRNAs with ATAC-seq from sCone photoreceptors (GSE83312). P-values and Odds Ratios from Fisher Exact Test. C) Venn diagram depicting the intersections of ATAC-seq from rods and ATAC-seq from sCones (GSE83312) with *Nrl*-repressed and *Nrl*-activated ncRNAs. P-values and Odds Ratios from Fisher Exact Test. D) Boxplots depicting the enrichment of Mean Reads per Million (RPM) of ATAC-seq from WT P21 mouse retinas and in *Nrl-/*-ATAC-seq P21 mouse retinas (left, GSE72550) and of Rod-specific ATAC-Seq and sCone specific ATAC-Seq (right, GSE83312) in *Nrl*-activated (green) and *Nrl*-repressed (blue) compared with background (Bkg) non-Nrl dependent ncRNAs (red). P-value calculated by ANOVA testing. E) Scatterplot depicting correlation of differentially expressed ncRNA transcripts with RPM of differentially expressed ATAC regions from rod (red) and cone photoreceptors (blue). Log2 (sCone ATAC RPM/Rod ATAC RPM) vs log2(*Nrl*^-/-^ ncRNA RPM/*Nrl*^+/+^ ncRNA RPM). Non-significantly changed transcripts in gray. F) Venn diagram depicting the overlap of annotated rod and cone specific elements as defined by high density reads^43,44,55^ with *Nrl*-activated ncRNAs (left) and *Nrl*-repressed ncRNAs (right). P-values and Odds Ratios from Fisher Exact Test.

We asked if *Nrl*-activated and *Nrl*-repressed ncRNA expression levels were concordant with the predicted strength of rod vs. cone CREs, as defined by the local quantitative enrichment of ATAC-seq read density (Figure 2D). We compared *Nrl*-activated and *Nrl*-repressed ncRNAs to wild-type and *Nrl*-deficient ATAC-seq reads (Figure 2D, left). We observed that CREs with *Nrl*-activated ncRNAs were associated with significantly higher ATAC-seq read density in the WT retina compared with the *Nrl*^-/-^ retina (Figure 2D; p=3.1e-5) (GSE72550)^19^. Conversely, we observed that CREs with *Nrl*-repressed ncRNAs were associated with significantly higher ATAC-seq read density in the *Nrl*^-/-^ retina compared with the WT retina (Figure 2D, p= 4.5e-10). (GSE72550)^19^.

We next assessed the association of *Nrl*-activated and *Nrl*-repressed ncRNAs with ATAC-seq read density in sorted rods and cones (GSE83312). We found that *Nrl*-activated ncRNAs were enriched at regions of higher ATAC-seq signal in rods than in cones (Figure 2D; p= 2.3e-5). In contrast, *Nrl*-repressed ncRNAs were enriched at regions of higher ATAC-seq signal in cones than in rods (Figure 2D; p= 5.3e-11). These observations indicate that *Nrl*-dependent ncRNAs differentially associate with candidate rod versus cone CREs: *Nrl*-activated ncRNAs affiliated with strong rod ATAC-seq signal, whereas *Nrl*-repressed ncRNAs affiliated with strong cone ATAC signal.

We hypothesized that the *Nrl*-dependent change in ncRNA may correlate with cell type specific rod versus cone change in chromatin accessibility. We therefore compared the *Nrl*-dependent change in ncRNA transcription at rod- and cone-specific ATAC-seq regions to the relative ATAC-seq read enrichment in rods vs. cones at those locations. We observed a positive correlation between downregulated ncRNAs and rod ATAC-seq reads, and between upregulated ncRNAs and cone ATAC-seq reads (Figure 2E, Cor=0.6, P<2.2e-16). This result indicated that the direction and quantitative degree of *Nrl*-dependence of ncRNA expression correlated with the relative cell-type specificity of the ATAC-seq signal, for *Nrl*-activated ncRNAs with rod-specific ATAC-seq and *Nrl*-repressed ncRNAs with cone-specific ATAC-seq.

We compared regions with high-density putative regulatory elements^43,44^ from rod and cone ATAC-seq with the *Nrl*-activated and *Nrl*-repressed ncRNAs. We observed that *Nrl*-activated ncRNAs predicted clusters of putative rod enhancers (Figure 2F, OR = 1.7, p = 4.7e-7) but not cone (Figure 2F, OR 1.05, p = 0.64). In contrast, *Nrl*-repressed ncRNAs predicted high density regulatory elements^43,44^ from cone (Figure 2F, OR = 1.6, p = 1.9e-4) but not rod cells (Figure 2F, OR = 0.95, p = 0.87). We conclude that *Nrl*-activated and repressed ncRNAs enrich two distinct regulatory pathways, identifying strong candidate CREs specific to rod and cone photoreceptors, respectively.

Mouse photoreceptor CREs are highly enriched for binding sites of CRX, a homeobox TF required for both rod and cone gene expression^14,17,42^. We asked if CREs defined by *Nrl*-dependent ncRNAs were CRX-bound. We observed significant enrichment for CRX occupancy by ChIP-seq in wild-type whole retinal tissue at *Nrl*-dependent ncRNA-defined regions of accessible chromatin (Figure 3A, top, GSE20012)^15^. Interestingly, using ATAC-seq data in conjunction with ncRNA transcription did not improve identification of CRX-bound elements (Figure 3A, bottom), suggesting that differential ncRNA transcription independently identifies CREs likely to be bound by CRX, without the need for CRX ChIP. Regions characterized as either *Nrl*-activated or repressed from the ncRNA analysis of the *Nrl*^-/-^ retina identified genomic CRX localization in wild-type tissue (Figure 3B, left). In *Nrl*^-/-^ retina, only *Nrl*-repressed ncRNAs identified CRX localization, providing candidates for cone-specific CRX occupancy, as expected with the absence of rod fate in the *Nrl*^-/-^ retina, and consistent with analysis of CRX occupancy in *Nrl*^-/-^ retinas (Figure 3B, right)^15^.

**Figure 3:**
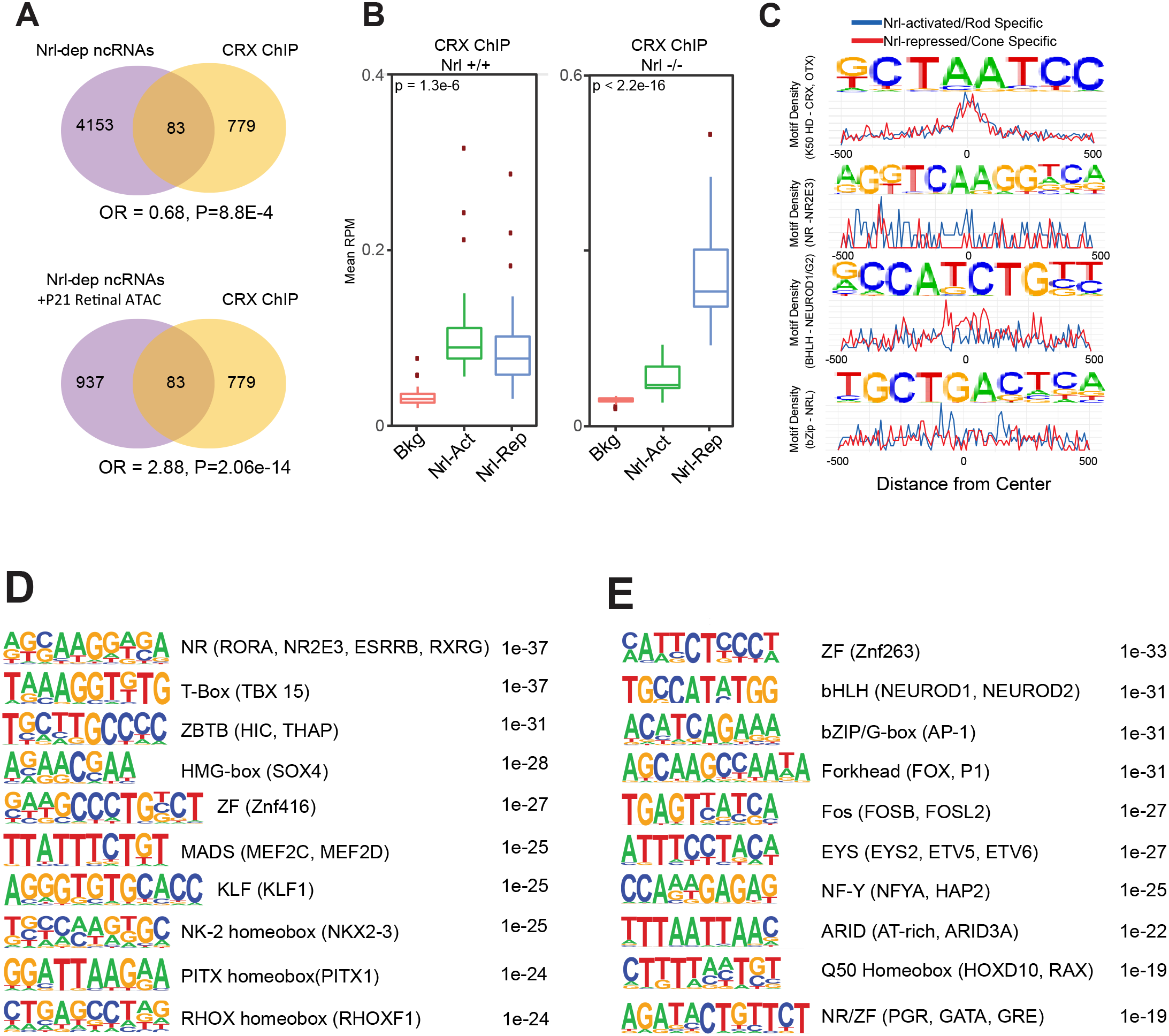
ncRNA-defined cell-type specific regulatory elements are enriched for cell-type specific TF motifs. A) Venn diagram of the overlap of total *Nrl*-dependent ncRNAs (top) and the intersection of *Nrl*-dependent ncRNAs with Retinal ATAC (GSE74661, bottom) with CRX bound sites from ChIPseq in whole mouse retinas (GSE20012). B) Boxplots depicting the relative enrichment of *Nrl*-activated (green) and *Nrl*-repressed (blue) ncRNAs in control CRX ChIP retinas (left) and *Nrl*^-/-^ CRX ChIP retinas (right), in relation to the ncRNA background (red). P value calculated by ANOVA testing. C) Position Weight Matrix (PWM, top) of select *de novo* binding motifs identified by *Nrl*-dependent ncRNAs. Motif Density (bottom) as a function of distance from center (bp) in *Nrl*-activated (Blue) and *Nrl*-repressed (Red). Hbox motif (CRX, OTX), BHLH motif (NeuroD1, NeuroG2), NR (NR2E3) and bZip (NRL, Mafa) motifs shown. D) Position Weight Matrix (PWM) of rod-specific *de novo* binding motifs grouped by TF family and ranked by p-value. E) Position Weight Matrix (PWM) of cone-specific *de novo* binding motifs grouped by TF family and ranked by p-value.

We performed *de novo* motif analysis of *Nrl*-dependent ncRNA-defined CREs to identify potential transcriptional co-regulators. Both activated and repressed ncRNAs identified the K50 homeodomain motif (K50 indicating the presence of lysine at position 50 of the homeodomain), shared among known NRL co-regulators, CRX and OTX2^15,18,45^ (Figure 3C, row 1). We interrogated the motifs specific for *Nrl*-activated and *Nrl*-repressed CREs, attempting to identify motifs specific for rod or cone CREs, respectively. *Nrl*-activated CREs were significantly enriched for the NRL binding motif (bZip) and NR2E3 (NR), a direct downstream target of NRL^37^ (Figure 3C, rows 2 and 4). By contrast, *Nrl*-repressed ncRNAs were differentially enriched for NeuroD1 binding motif (BHLH, Figure 3C, row 3). Thus, *Nrl*-dependent ncRNAs identify *Nrl*activated and repressed CREs characterized by specific TF binding motifs.

To define potential novel transcriptional pathways that are unique to rod and cone photoreceptor cell types, we assessed comparative motif enrichment for *Nrl*-activated CREs in rod-specific open chromatin versus *Nrl*-repressed CREs in cone-specific open chromatin (Figure 3D-E). We identified TF motifs unique to rods (Figure 3D) and cones (Figure 3E). Consistent with comparative analysis in Figure 3C, we find the NR motif unique to rod-specific elements (Figure 3D), and the bHLH binding motif unique to cone-specific regulatory elements (Figure 3E) with this comparative analysis. These differential motifs predict rod and cone specific TFs that may help define the two cell types.

### *Nrl*-dependent ncRNAs define functional photoreceptor regulatory elements

We predicted that the ncRNA-defined candidate regulatory elements are active in photoreceptors, with at least a subset differentially active in rods versus cones. We first examined the regulatory capacity of *Nrl*-dependent ncRNA-defined CREs in the developing mouse retina. As *Otx2* is required for the genesis of rods and cones, the regulation of this locus is of great interest^25^. Data from the ENCODE project allowed us to identify 24 DNase I hypersensitivity sites (HS) at the *Otx2* locus, from mouse retina at P0, within 300kb around the *Otx2* gene (Figure 4A)^46^. These regions were compared with those identified through *Nrl*-dependent ncRNA profiling. Of the 24 ENCODE sites, 4 overlapped *Nrl*-repressed ncRNAs and cone ATAC-seq peaks (Figure 4B). We examined the activity of all 24 locations to determine if candidate CREs overlapping with *Nrl*-repressed ncRNAs would be distinguished from those without overlapping ncRNAs. The DNA sequences corresponding to the DNAse I HS sites were cloned into the reporter plasmid Stagia3, which has an eGFP-IRES-AP reporter^47,48^. We tested these constructs, along with a control, ubiquitously expressed CAG-Cherry plasmid, for activity in dissected mouse retina through electroporation at E14.5. E14.5 is a period of development when primarily cones, and not rods, are being generated, and electroporation into mature cones at later ages is inefficient^49^. Retinas were then cultured as explants on filters for 2 days. Nine out of the 24 DNAse I HS sites showed alkaline phosphatase (AP) activity (Figure 4A, C). Interestingly, three of the four ENCODE DNAse I HS sites corresponding with *Nrl*-dependent ncRNAs with P21 cone ATAC-seq peaks were among the most active, highlighting the observation that TF-dependent ncRNAs mark potent regulatory elements (Figure 4B).

**Figure 4:**
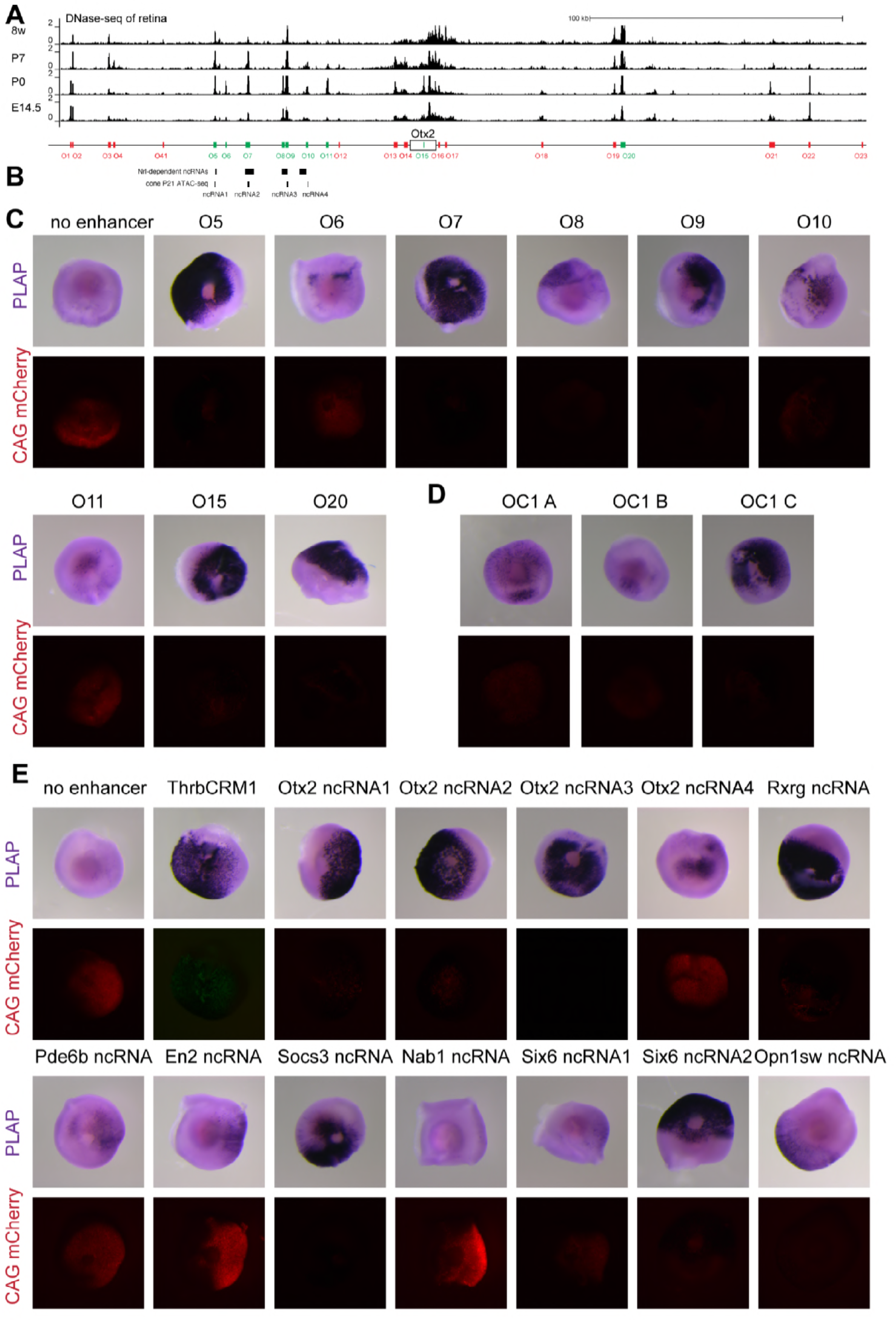
*Nrl*-repressed ncRNAs identify cone-specific regulatory elements with regulatory activity *ex vivo*. A) Annotation track shows ENCODE DNaseI Hypersensitivity Sites for mouse retinal cells at various stages of development for the Otx2 locus. 24 peaks were selected for further testing (O1-O23) within a ~300kb region centered around Otx2. B) *Nrl*-dependent ncRNAs at the Otx2 locus and the nearest ATAC-seq from cone P21 cells. C) DNA sequences corresponding to the DNAse I HS peaks annotated in (A) were cloned into the reporter plasmid Stagia3 (alkaline phosphatase (AP) and EGFP) and tested for activity in the mouse retina by ex vivo electroporation at E14.5 along with a co-electroporation control CAG-Cherry. Retinas were then cultured as explant on filters for 2 days. Positive regions are shown in green in (A), while the negative are in red. D) Electroporated constructs containing DNA sequences defined by retina DNAse I HS from ENCODE at the Onecut1 locus. E) Retinal explants electroportated with ncRNA DNA sequences defined by cone ATAC-seq peaks associated with the ncRNAs found at the Otx2 locus and other known retinal genes.

As cone development is regulated by both *Otx2* and Onecut 1 (*Oc1*), we also were interested in regulatory elements at the *Oc1* locus^25,50^. To this end, several regions predicted by the ENCODE DNAse I analysis were tested (Figure 4D). As with *Otx2*, a region with strong regulatory activity had an *Nrl*-dependent ncRNA. These results indicate that intersecting TF-dependent ncRNA expression with previously published ENCODE datasets provides novel information useful for identifying strong regulatory elements, compared to utilizing chromatin accessibility alone.

We hypothesized that *Nrl*-repressed ncRNAs may identify CREs with cone activity. To examine whether CREs defined by the overlap of *Nrl*-repressed ncRNAs and cone ATAC-seq peaks have activity in developing cones, we examined ncRNA defined elements from the *Otx2* and *Rxrg* loci, as these genes are known to be important for cone development^30,51^. The DNA sequences from these loci (material and methods) were cloned into the Stagia3 reporter plasmid. We also tested putative CREs defined by ncRNA expression for other genes involved in the development of the visual system (*En2, Socs3, Nab1, Sixβ, Opn1sw*), as well as *Pde6b*, a rod-specific gene. These plasmids were delivered by electroporation of ex-vivo E14.5 mouse retinas and assayed for AP activity 2 days later (Figure 4E). The ThrbCRM1-dtTomato construct was used as a positive control for AP, as it has known activity in cones, HCs, and a subset of RPCs that produce cones and HCs in the chick retina^25^. Ten of 12 candidate regulatory elements were able to drive AP expression in explanted E14.5 mouse retinal tissue (Figure 4E).

To define the specific cell-types with reporter activity, active enhancer constructs were tested for expression of eGFP, which provided greater cellular resolution and coincident assay with cell-type specific marker co-expression. The ThrbCRM1-tdTomato plasmid was first tested for its use as a positive control to specifically mark murine cones. ThrbCRM1-tdTomato positive cells were located in the apical region of electroporated E15.5 retinas (Figure 5A), which is the location of developing cones^18,25^, and they were positive for the Rxrg protein, a validated cone marker (Figure 5B). We then examined co-expression of eGFP, driven by the *Nrl*-repressed ncRNA constructs, and tdTomato from ThrbCRM1 following electroporation into E14.5 retinal explants. GFP-positive cells from 7 of the *Nrl*-repressed ncRNA constructs were located in the apical region and showed a strong overlap with tdTomato (Figure 5C). The only exception was the *Socs3* element (Figure 5C), which marked cells with a morphology and position matching those of RPCs. We further tested the expression from the *Otx2* ncRNA-defined enhancer regions 1 and 2 with that of the OTX2 protein (Figure 5D). Both Otx2-ncRNA-defined elements showed strong co-localization with the Otx2 protein. Together, these findings indicate that CREs marked by *Nrl*-dependent ncRNAs in the *Nrl* mutant setting have activity in developing cones.

**Figure 5:**
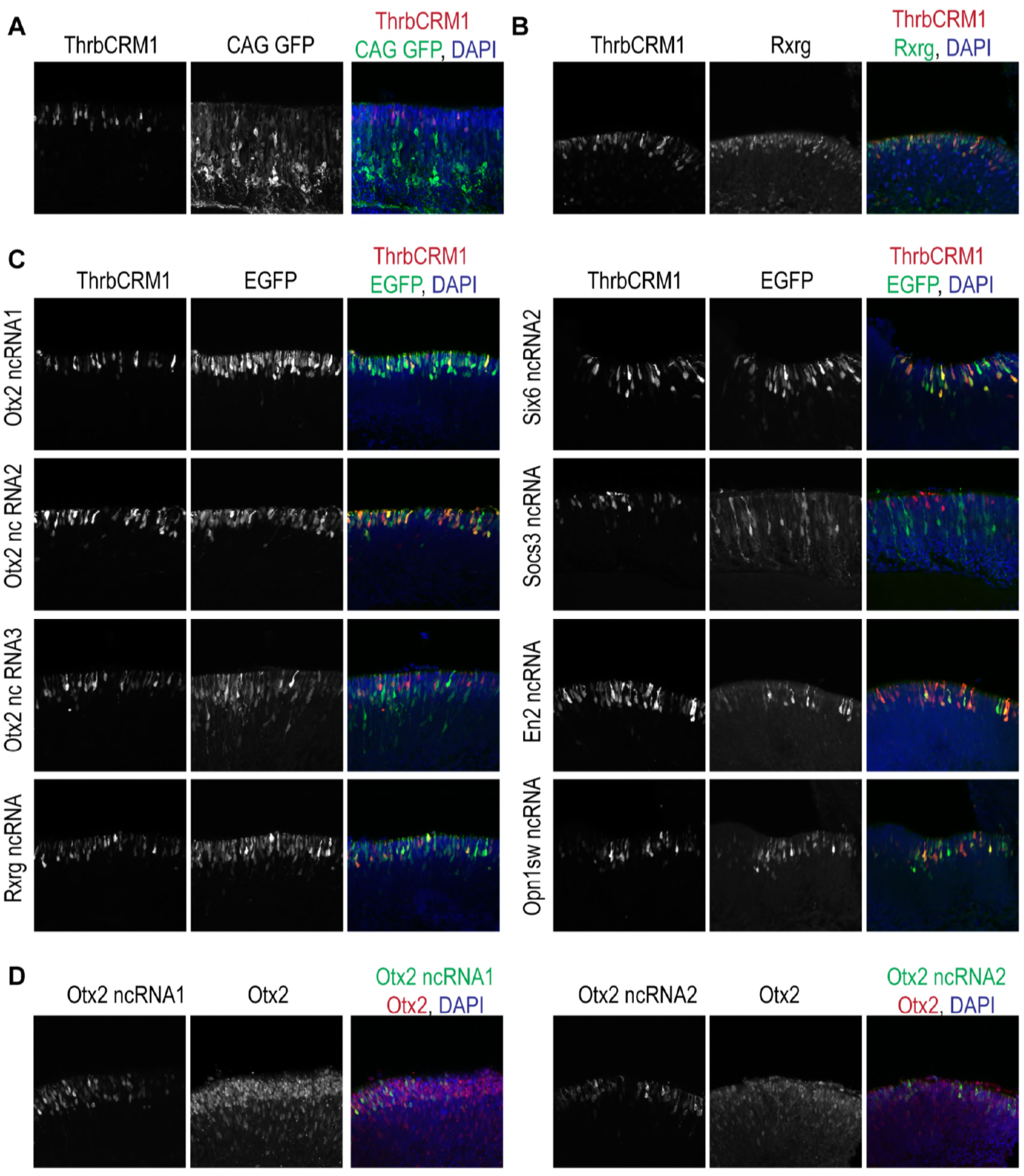
*Nrl*-repressed ncRNAs identify functional cone-specific regulatory elements *in vivo*. A) Transversal sections of retina from mice electroporated at E15.5 (A) or E14.5 (B-D) and cultured for 2 days before fixation and embedding are shown. The ThrbCRM1-dtTomato enhancer was electroporated into mouse retina with an ubiquitous CAG-GFP co-electroporation control. ThrbCRM1 positive cells are located in the apical region of the developing ONL, where photoreceptors are found (left). The co-electroporation GFP reporter is expressed in mitotic cells at the time of the electroporation. B) The ThrbCRM1 enhancer (left) was expressed in cells positive for Rxrg protein (center), a cone marker gene. C) Positive regions from our AP screen were co-electroporated (Figure 5 E) along with the ThrbCRM1-dtTomato enhancer (left). The Stagia3 reporter plasmid used for AP also contains an eGFP readout (center column). All but one of the cells positive for the putative enhancers tested are located in the developing ONL, and show a strong overlap with the cone marker ThrbCRM1. D) Two Nrl-dependent ncRNAs at the Otx2 locus were tested (left column) for their co-localization with the Otx2 protein (center). Both showed a strong overlap (composite image, right column).

## Discussion

### Application of enhancer transcription to the definition of *cis*-Regulatory Elements

Defining the GRNs that distinguish healthy and disease states requires the identification of the functionally relevant CREs and TFs that regulate them. Current approaches for nominating CREs, including histone modifications, chromatin status^3–5^ and TF occupancy^6^, have been successful in many contexts. However, only a small fraction of the thousands of candidate TF-dependent enhancers identified by these approaches have been functionally validated^52^. Moreover, these approaches do not reveal context specificity of expression from a region, nor provide a quantitative assessment of enhancer function^1,53^. We previously defined a complementary approach to regulatory region identification, utilizing context-dependent enhancer transcriptional profiling, to nominate regulatory elements. This approach defined a GRN composed of functional elements in cardiomyocytes defined by ncRNA expression from the adult mouse heart that contribute to cardiac rhythm control^1^. Here we extended TF-dependent ncRNA profiling to define regulatory regions that govern photoreceptor gene expression in the retina and assess the applicability of this approach for defining both wild-type and mutant elements that comprise context-specific GRNs.

### Regulatory regions active in photoreceptors and their progenitor cells

An understanding of the GRNs that govern photoreceptor production is particularly important, given the desire to generate cones from stem cells for therapeutic applications^13^. Definition of the GRN for cone genesis has been limited, as cones are typically born early, when access is limited, and cones are much less abundant than rods. The conversion of rods to cone-like cells in the *Nrl*^-/-^ mouse has provided a deeper understanding of the molecular components of both cones and rods, including the identification of rod- and cone-enriched transcripts^18,28^, as well as the delineation of rod and cone open chromatin regions^18,19^. These datasets have provided an excellent background for an assessment of TF-dependent ncRNA profiling as a method for nominating regulatory regions near *Nrl*-dependent genes. Consistent with a correlation between ncRNA transcription and regulatory region activity, the *Nrl*-dependent ncRNAs defined regions that had a high density of marks of regulatory regions, such as those proximal to *Rxrg, Otx2*, and Gasβ^7–10, 43,44^

*Nrl*-dependent ncRNAs that were upregulated in the *Nrl* mutant retina identified candidate regulatory regions associated with cone genes, and ncRNAs that were downregulated in the *Nrl* mutant identified candidate regulatory regions associated with rod genes. Functional examination of enhancer activity for a subset of these predicted elements at relevant cone genes showed cell type specificity for the majority of *Nrl*-repressed elements (Figure 5). However, a small number of tested elements did not display the predicted cone pattern of activity. A few had activity in other cell types, or had no specific activity in retinal explants. Some DNase I hypersensitivity sites from the Otx2 locus where we found *Nrl*-repressed elements also had activity in bipolar interneurons (not shown, Wang et al. in preparation), which are born in the postnatal period, and in rods^54^. This may be due to the fact that the CREs defined by ncRNAs and ATAC-seq peaks are typically fairly large and thus may harbor multiple TF binding sites. These binding sites may be rod, cone, or bipolar cell-specific, and may rely on higher ordered chromatin structure for proper regulation, structure that likely is not included in electroporated plasmids. Elements that showed no specific activity in retinal explants may be active in mature cones, and not in the cone progenitor cells or immature cones that were assayed here.

While many of the genes associated with the ncRNA-repressed list were related to cone development or function (*Otx2, Rxrg, Gngt2, Gnat2, Opn1sw, Sall3*)^30,51,55–57^, we also found genes that are generally important for eye or retina development that have not been well characterized with respect to cone-specific development. Interestingly, the ncRNA-defined CRE associated with *Six6*, an eye-field TF, or the element near *En2*, a gene important for ganglion cell differentiation^18,58^, displayed reporter activity in cones. These results suggest that non-coding transcriptional profiling uncovered not only rod- and cone-specific regulatory programs, but potential shared regulatory programs that warrant further investigation.

### Defining TF-dependent networks of cell fate has implications for the study of disease-specific regulatory pathways

Defining both activated and repressed regulatory programs has significance across biological contexts for distinguishing the CREs driving gene regulation in normal versus mutant, or disease, states. Disease GRNs have been primarily characterized via discovery of wild-type enhancers, followed by assessment of disease-specific changes in their activity. This presumes a model in which disruption of the wild-type GRN sufficiently describes the disease state. However, our findings highlight an emergent, mutant-specific, GRN through the use of context-specific CRE activity. Defining regulatory pathways in NRL presence and absence parallels potential studies of healthy versus diseased states. Defining such context-specific CREs in disease states may define novel regulatory pathways that mediate disease pathogenesis, with potential therapeutic implications. The effectiveness of enhancer transcriptional profiling for identifying emergent networks suggests a potential for future application, for the assessment of both cell-type specific and disease GRNs.

## Acknowledgments

This research was supported in part by NIH R01 HL092153, R01HL124836, and R33 HL123857 to IPM, R01EY024958, R01EY025196, and R01EY026672 to JCC, F30 HL131298 to RDN, AHA Collaborative Sciences Award to IPM, and HHMI (CLC). This research was supported in part by the Leducq Foundation (IPM). This research was supported in part by the NIH through resources provided by the Computation Institute and the Biological Sciences Division of the University of Chicago and Argonne National Laboratory, under grant 1S10OD018495-01. N.L. was supported by post-doctoral fellowships from the Swiss National Science Foundation and the Human Frontiers Science Program.

## Materials and Methods

### Animals

*Nrl*^-/-^ mice were generated as previously described^18,59^. Mouse husbandry and all procedures (including euthanasia by CO2 inhalation and cervical dislocation) were conducted in accordance with the Guide for the Care and Use of Laboratory Animals of the National Institutes of Health, and were approved by the Washington University in St. Louis Institutional Animal Care and Use Committee. For ex-vivo enhancer testing, wild-type embryos were obtained from timed pregnant CD1 mice (Charles River Laboratories). All animal studies were approved by the Institutional Animal Care and Use Committee at Harvard University.

### Coding RNA-Seq library preparation and Data analysis

Libraries were prepared from this RNA starting with 1 g per sample and using the mRNA-seq Sample Prep Kit (Illumina) as per recommended instructions. After Ribozero purification and removing only ribosomal RNA, barcoded libraries were prepared according to Illumina’s instructions (2013) accompanying the TruSeq RNA Sample prep kit v2 (Part# RS-122-2001). Libraries were quantitated using the Agilent Bio-analyzer (model 2100) and pooled in equimolar amounts. The pooled libraries were sequenced with stranded 50-bp single-end reads on the HiSeq2500 in Rapid Run Mode following the manufacturer’s protocols (2013).

RNA library preparation was performed as previously discussed^1^. Briefly, 22M to 30M reads were mapped to mouse genome with TopHat2 (v 2.1.1). Reads mapped to the mitochondrial genome, and with phred score < 30 were excluded. Counts were retrieved with HTseq (v.0.6.0)^60^ in union mode. Lastly, counts were analyzed for differential expression with R (3.4) package DEseq2^61^.

### Noncoding RNA-Seq library preparation

Total RNA was extracted by TRIzol Reagent (Invitrogen), followed by ribosomal and polyA depletion. After RiboZero purification and oligo-dT depletion, RNA Barcoded Libraries were prepared according to Illumina’s instructions (2013) accompanying the TruSeQ RNA Sample prep kit v2 (Part# RS-122-2001). Libraries were quantitated using the Agilent Bio-analyzer (model 2100) and pooled in equimolar amounts. The pooled libraries were sequenced with 50-bp stranded single-end reads on the HiSEQ4000 in Rapid Run Mode following the manufacturer’s protocols (2013).

### Noncoding RNA-Seq Data analysis

About 170-186 million high-quality reads (quality score >30) reads for each sample were obtained. Fastq files were aligned to UCSC genome build mm9 using TopHat (version 2.0.10) as previously described^62^ and between 168 million and 174 million reads were successfully mapped. RABT assembly was performed by Cufflinks (version 2.2.1, with parameters -g -frag-bias-correct -multi-read-correct -upper-quartile-norm), as it can recover transcripts that are transcribed from segments of the genome that are missing from the current genome assembly. Analysis of differential expression was performed using Cuffdiff from the Cufflinks package^62^. False discovery rate (FDR) was calculated after removing the coding-gene counts. Significance was considered to have been reached when FDR was <0.05 and fold change was > 2. The mm9 genomic coordinates of identified noncoding transcripts were lifted over to the mouse mm10 before comparisons with open chromatin and TF-binding regions.

### GO enrichment analysis

Enrichment of GO Biological Process terms from genes within 2MB of ncRNAs was performed with Bioconductor package GOstats version 2.46^63^.

### ATAC-seq and ChIP-seq data processing

Fastq files from previously generated ChIP-seq (GSE20012) and ATAC-seq (GSE83312) datasets were downloaded from GEO and processed identically as previously described^18^. Briefly, adapter sequences were clipped from reads using Cutadapt^64^, then aligned to UCSC mouse genome mm10 with bowtie version 2.3.4^65^ in end-to-end mode. Mismatched reads, PCR duplicates, ENCODE blacklisted regions, and reads with quality < 30 were removed with Samtools version 1.5^66^. For ATAC-seq, fragments with width > 147 base pairs were removed to enrich for nucleosome free reads using a custom script. Peaks for both assays were called with Macs 2.11^67^.

### Associating ncRNAs and regulatory elements

Open chromatin peaks and TF-binding peaks were intersected with ncRNAs using Bioconductor package GenomicRanges^54^ allowing for a 500bp gap.

### Metagene analysis

To compare the coverage of ChIP-seq and ATAC-seq regions in differentially expressed ncRNAs we used Bioconductor Package Metagene (2.12.1). Coverage was normalized to reads per million. We binned the position of each region to 100 bp. We modified the current Metagene source code to output boxplots as opposed to ribbons and we then tested the difference of means with ANOVA.

### Identification of differentia//y accessible peaks

Rod and cone ATAC-seq peak regions were combined and sorted with bash commands (cat sort). Counts were retrieved from each alignment file using Bedtools multicov (2.26.0) and tested for differential expression with DESeq2^. Peaks were considered differentially accessible when log2 fold change greater than 1 and p-adjusted value less than 0.05 and these regions were considered cone-specific and rod-specific. To assess the relationship between differentially expressed ncRNAs and differentially accessible open chromatin we performed a global overlap of all ncRNA regions and combined ATAC-seq regions. Then differential regions from both sets were highlighted.

### Motif Enrichment analysis

Known and de novo motif scanning was performed with HOMER (4.3) using Rod-specific and cone-specific ATAC-seq regions that intersected with *Nrl* wildtype ncRNAs and *Nrl* null ncRNAs respectively. Target sequences consisted of 200 bp elements centered on peak summits. Background sequences consisted of approximately 50,000 randomly selected 200 bp intervals from the mouse genome normalized for mono- and di-nucleotide content relative to each target set. Repeat sequences were masked from the genome, and targets with >70% of bases masked were dropped from enrichment analysis. Preferential spacing between highly enriched motifs (MAF, bZIP, and NR motifs) in differential regions was assessed by first centering the above intersect on individual motifs and plotting the density.

### Identification of high density peaks regions

Cone and Rod specific high density peak regions were marked by HOMER (4.3) using a 12.5kb window^43,44^. Alignment files for rod and cone ATAC-seq were used to find peaks with Homer using the parameter -style Super.

### Retina electroporation and AP staining

Ex vivo retina electroporation was carried out as described previously^68,69^, with at least three biological replicates for AP staining, or at least in duplicates for immunohistochemistry. The chamber used for electroporation was modified as previously described^70^. Stages of embryos used for the experiments are described in the main text of in the figure legends. Electroporation settings were 5×30 V pulses, 50 ms each and 950 ms apart. DNA concentration was 400-600ng/ul for control plasmids and 1ug/ul for enhancer constructs. Retinas were harvested after 2 days in culture.

### Plasmid and DNA sequences

In vivo enhancer testing was performed with the Stagia3 reporter vector (Addgene #28177)^48^. Enhancer testing with the CAG-EGFP, CAG-mCherry, and ThrbCRM1-tdTomato vectors were modified from our previous work^25,69^. Coordinated of regions cloned are shown in mm10 assembly.

O10> chr14:48616314-48617497

O11> chr14: 48624486-48626389

O5> chr14: 48579937-48581029

O6> chr14: 48584564-48585012

O7> chr14: 48593170-48594188

O8> chr14: 48606973-48608016

O9> chr14: 48608144-48609697

O15> chr14:48662841 -48663211

O20> chr14:48740203-48742409

Oc1 A> chr9:74384085-74384740

Oc1 B> chr9 74530189-74532399

Oc1 C> chr9:74810971 -74812406

Otx2 ncRNA1>chr14 + 48580418 48580655

Otx2 ncRNA2>chr14 + 48593310 48594038

Otx2 ncRNA3>chr14 + 48608844 48609310

Otx2 ncRNA4>chr14 + 48617045 48617321

Rxrg ncRNA>chr1 + 167438156 167438645

Pde6b ncRNA>chr5 + 108366779 108367075

En2 ncRNA>chr5 + 28145373 28145644

Socs3 ncRNA>chr11 + 117963224 117963787

Nab1 ncRNA>chr1 + 52435750 52436194

Six6 ncRNA1>chr12 + 72854405 72854809

Six6 ncRNA2>chr12 + 72831804 72832279

Opn1sw ncRNA>chr6 + 29394311 29394823

### IHC

20-30um retinal section were prepared and stained as described previously^68^. Blocking solution was: 0.3% Triton in 1× PBS. Primary antibodies used in this study include: chicken anti-GFP (1:1000, Abcam, AB13970), rabbit anti-mCherry (1:1000, Abcam, 167453), rabbit anti-Otx2 (1:500, Millipore, AB9566), mouse anti-Rxrg (1:300, Santa Cruz Biotechnology, sc-514134). Secondary antibodies were from Jackson Immunoresearch.

### Imaging

Retina explants were imaged on a Leica M165FC microscope. Retinal section images were acquired by a Zeiss LSM780 inverted confocal microscope from the Microscopy Resources on the North Quad (MicRoN) core at Harvard Medical School.

---

**Figure S1.**
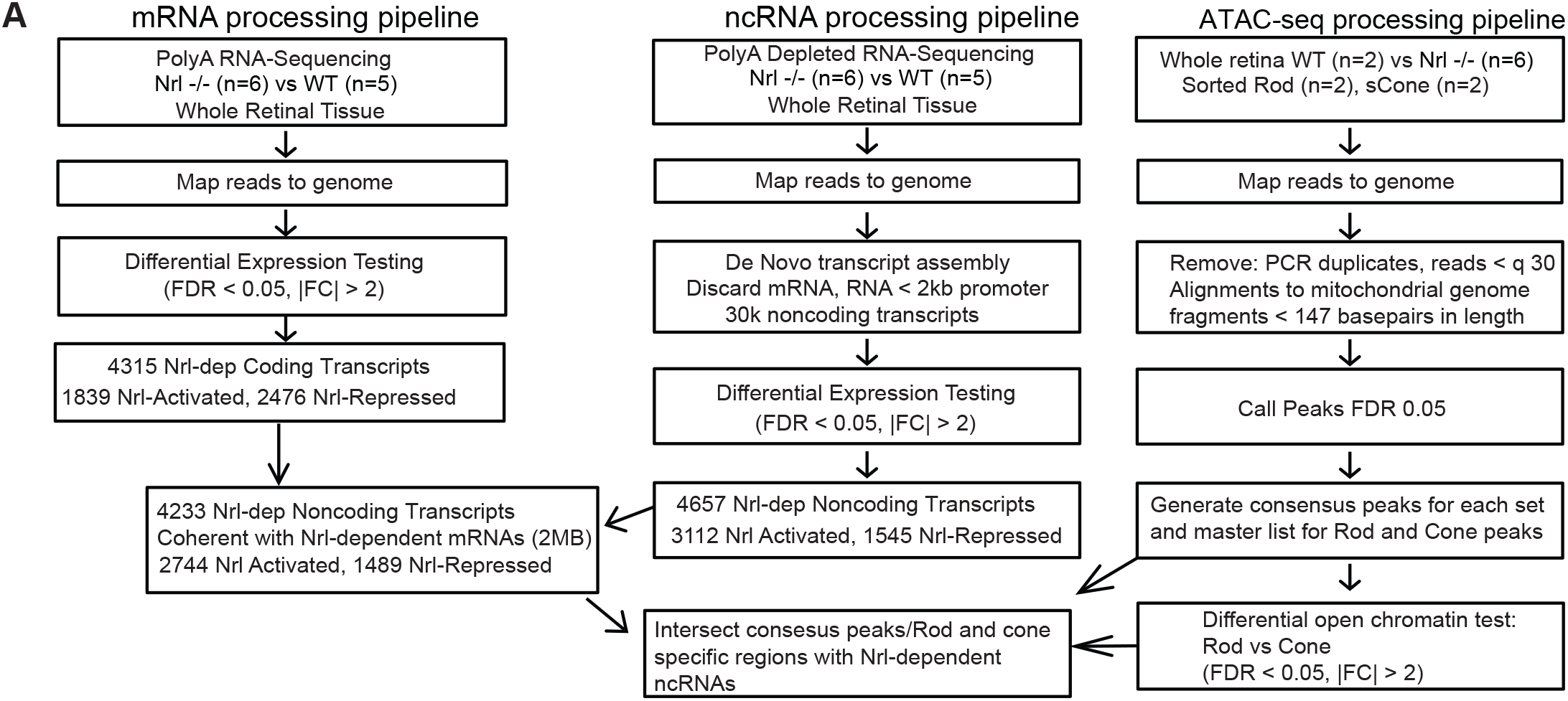
Processing pipelines for mRNA-seq, ncRNA-seq and ATAC-seq.

